# Identification of large offspring syndrome during pregnancy through ultrasonography and maternal blood transcriptome analyses

**DOI:** 10.1101/2022.03.31.486611

**Authors:** Rocío Melissa Rivera, Anna Katherine Goldkamp, Bhaumik Narendrabhai Patel, Darren Erich Hagen, Edgar Joel Soto-Moreno, Yahan Li, Chris Kim, Cliff Miller, Fred Williams, Elizabeth Jannaman, Yao Xiao, Paula Tribulo, Eliab Estrada-Cortés, Astrid Roshealy Brau-Rodríguez, Peter James Hansen, Zhoulin Wu, Christine Marie Spinka, Neal Martin, Christine G. Elsik

**Affiliations:** Division of Animal Sciences, University of Missouri, Columbia, MO; Department of Animal and Food Sciences, Oklahoma State University; Green Hills Veterinary Clinic, MO; Department of Veterinary Pathobiology, University of Missouri, Columbia, MO; Department of Animal Sciences, University of Florida, Gainesville, FL; Department of Health Management and Informatics, University of Missouri, Columbia, MO; Martin Veterinary Services, Centralia, MO

**Keywords:** Large offspring syndrome, abnormal offspring syndrome, assisted reproductive technologies, fetal morphometry, maternal blood biomarker, Beckwith-Wiedemann Syndrome

## Abstract

The use of assisted reproductive technologies (ART) in cattle can result in large/abnormal offspring syndrome (LOS/AOS) which is characterized by macrosomia. LOS can cause dystocia and lead to the death of dam and calf. Currently, no test exists to identify LOS pregnancies. We hypothesized that fetal ultrasonography and/or maternal blood markers are useful to identify LOS. Bovine fetuses were generated by artificial insemination (control) or ART. Fetal ultrasonographies were taken on gestation day 55 (D55) and fetal collections performed on D56 or D105 (gestation in cattle ≈280 days). ART fetuses weighing ≥97 percentile of the control weight were considered LOS. Ultrasonography results show that the product of six D55 measurements can be used to identify extreme cases of LOS. To determine whether maternal blood can be used to identify LOS, leukocyte mRNA from 23 females was sequenced. Unsupervised hierarchical clustering grouped the transcriptomes of the two females carrying the two largest LOS fetuses. Comparison of the leukocyte transcriptomes of these two females to the transcriptome of all other females identified several misregulated transcripts on gestation D55 and D105 with *LOC783838* and *PCDH1* being misregulated at both time-points. Together our data suggest that LOS is identifiable during pregnancy in cattle.

## INTRODUCTION

Large offspring syndrome (LOS) is an overgrowth condition observed in ruminant fetuses and neonates^1,2^. LOS was first reported in 1991 in a cloned calf produced via nuclear transfer^3^. Later, in 1995, overgrowth was reported as a result of non-invasive assisted reproductive technology (ART) procedures^4^. At that time, the overgrown animals were called “large calves” and the syndrome was coined LOS^5^. As the use of ART in ruminants increased during the next decade, so did the number of LOS reports^6-11^. The main characteristic of LOS is overgrowth, which in some instances can result in calves weighing twice the average birthweight of their breed^4^. However, LOS in ruminants is a complex disorder with other phenotypes observed including visceromegaly, macroglossia, increased incidence of hydro-allantois, abnormal limbs and spinal cord, ear malformation, hypoglycemia, and umbilical hernia^6,7,9-14^. Because of these varied phenotypes, this syndrome is also known as abnormal offspring syndrome (AOS)^11^.

Even though it is now clear that LOS/AOS is a multi-locus loss-of-imprinting (i.e., epigenetic) condition^14^, it is still not known what triggers LOS and which ART (e.g., *in vitro* maturation of oocytes, *in vitro* fertilization, *in vitro* embryo culture or embryo transfer) is involved. Several reported cases of LOS in the literature were produced using serum supplementation during oocyte maturation and/or during embryo culture, which suggest that serum may be a factor promoting the syndrome^1,2,4,15,16^. Serum has been experimentally determined to cause LOS in sheep^16-18^ and bovine offspring derived from embryos cultured in serum containing medium can develop LOS ^2^. In addition, the syndrome can also occur in fetuses and calves derived from embryos cultured without serum supplementation^19,20^ and, more recently, we have documented that this syndrome occurs spontaneously in cattle produced by natural or artificial insemination ^21-23^. The latter is of interest as there is a similar loss-of-imprinting overgrowth syndrome in humans, namely Beckwith-Wiedemann syndrome, which occurs naturally, and its incidence is increased in children conceived by ART ^24^.

Due to its large size, LOS can cause dystocia and, sometimes, cesarean section is needed for delivery ^25^. Even if the newborn calf survives the difficult birth, the enlarged tongue or extreme body weight make suckling difficult, thus increasing the chances of postnatal death ^26^. In addition to the possible death of calves and cows, other financial losses are incurred due to veterinary costs ^22^ and the associated negative economic impact in terms of losses in milk, fat, and protein yields in the subsequent lactation ^27,28^. For example, two independent LOS cases have been recently reported with total estimated losses of approximately $30,000 each^22^. These monetary losses could have been minimized if the early identification of LOS was possible. To date, however, no test exists to predict LOS pregnancies in cattle. As ART is the current method of choice to improve genetic merit of the offspring in the cattle industry ^29,30^ it is of particular importance to find biomarkers to identify fetal overgrowth early during gestation to help producers decide whether to terminate the pregnancy or prepare for a difficult birth.

Ultrasound is a valuable non-invasive and repeatable tool that has been widely used in cattle to determine fetal growth^7,31,32^, fetal age^33^, fetal sex^34^ and clinical pathologies such as mummified fetuses or endometritis^35^. In addition, blood biomarkers have been successfully used as a non-invasive method to determine pregnancy status. For example, *ISG-15* mRNA from pregnant cattle leukocytes^36^ and pregnancy associated glycoproteins from bovine maternal blood serum are useful markers of early pregnancy^37^. Whether maternal blood components can be used to identify LOS early in pregnancy is not known.

For our study, we hypothesized that LOS can be identified during pregnancy in cattle by use of ultrasonography and/or maternal blood leukocyte mRNA biomarkers. The approach was to generate embryos by artificial insemination (AI; control) or by *in vitro* procedures previously shown by us to generate overgrown fetuses and to perform ultrasonographic measurements of those fetuses at day 55 of gestation (gestation length in cattle ≈ 280 days) and transcriptome analysis of maternal blood leukocytes on days 55 and 105 of gestation.

## MATERIALS AND METHODS

### Heifers

All animal procedures were conducted in accordance with the Guide for the Care and Use of Agricultural Animals in Research and Teaching and approved by the Institutional Animal Care and Use Committee of the University of Missouri (Protocol #9455). All animals were kept at the University of Missouri South Farm Research Center in Columbia.

Angus crossbred heifers of approximately 18-20 months of age were synchronized and selected for breeding using the 14-day controlled internal drug release (CIDR®, Zoetis, Kalamazoo, MI) prostaglandin and timed artificial insemination protocol (**Supplemental Figure 1**).

**Figure 1.**
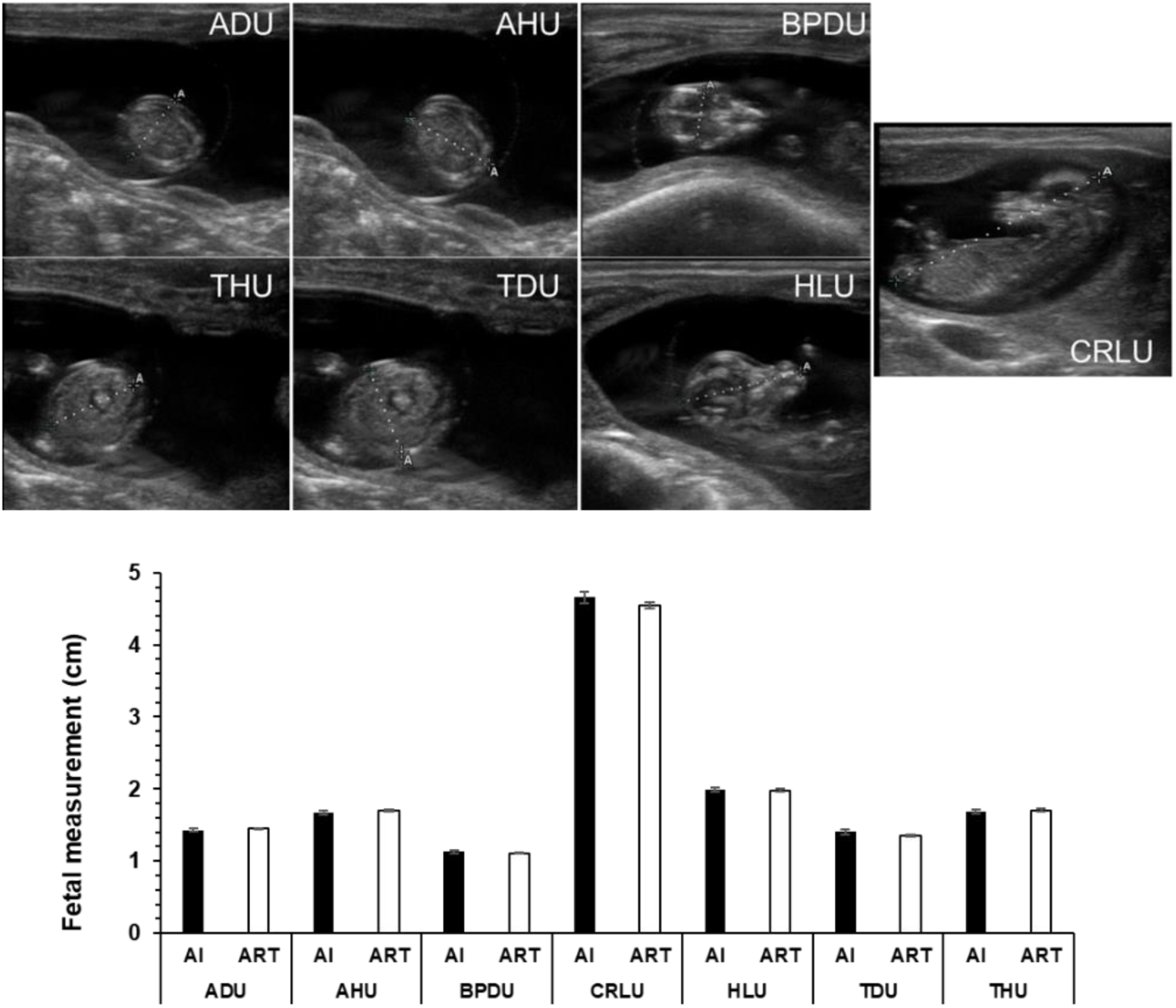
Fetal morphometry of fetuses collected on day 105 gestation. **Top**. sagittal views of the ultrasound images taken on day 55 of gestation. U: ultrasound. AH: abdominal height. AD: abdominal diameter. BPD: biparietal diameter. CRL: crown rump length. HL: head length. TD: thoracic diameter. TH: thoracic height. **Bottom:** Comparison of ultrasound measurements between all the AI (n=26) and ART (n=89) fetuses in this study. No statistical differences were detected between groups. AI: artificial insemination. ART: assisted reproductive technology (*i*.*e. in vitro* produced embryos).

### *In vitro* and *in vivo* production of embryos and embryo transfer

Media and procedures were as previously described by us^2,38^. Briefly, *Bos taurus taurus* (*B. t. taurus;* Angus/Angus-Crossbred) oocytes were harvested from slaughterhouse ovaries. Oocytes were removed from maturation medium after ∼21 h of culture and inseminated with semen from one *B. t. indicus* male (Brahman breed [JDH MR MANSO 7 960958 154BR599 11200 EBS/INC CSS 2]). Putative zygotes were stripped of cumulus cells by five minutes of vigorous vortexing at approximately 18 h after insemination and then cultured in KSOM supplemented with amino acids in a humidified atmosphere containing 5% O_2_, 5% CO_2_, and 90% N_2_. On day five after insemination, the culture medium was supplemented with 10% (v/v) estrus cow serum (collected and prepared in house and previously used in ^2^) and embryos returned to the incubator. On day seven, blastocyst-stage IVP embryos were selected, washed in BioLife Holding & Transfer Medium (AgTech; Manhattan, KS), and loaded in groups of two into 0.25 cc yellow, direct transfer and irradiated straws (AgTech). Blastocysts were transferred to synchronized recipient females on day seven after estrus (**Supplemental Figure 1**).

### Rationale for experimental design

This experiment is a part of a large-scale study aimed at the identification of epigenetic misregulations in LOS (as defined by ART fetuses weighing ≥97 percentile of the weight of the control fetuses). Based on our previous results for pregnancy rates from IVP embryos, percent LOS on D105, and transcriptome and methylome analyses^39^, we calculated that four LOS males and four LOS females would be required to achieve 90% power. Therefore, we transferred two embryos per heifer (as we did in^2^) to achieve that goal. In our previous study ^2^, body weight did not differ between singletons and twins on D105 of gestation. It should be noted that twinning occurs in cattle^40^. In addition, the rationale for generating *Bos taurus indicus* x *Bos taurus taurus* F1 fetuses in this study is that since LOS is a loss-of-imprinting condition, allele sequence differences are required to characterize parental-specific genomic (mis)regulation in the F1. This has been a useful breeding scheme to enhance the list of imprinted genes in bovine and to identify loss-of-imprinting in LOS^39,41^.

### D55 and D77 fetal ultrasound morphometry

At day 55 (D55) of pregnancy, presumed pregnant animals were checked for the presence of a fetus/es by transrectal ultrasonography using a SonoSite EDGE ultrasound machine equipped with a L52 10.0-5.0 megahertz linear-array transducer (**Supplemental Figure 1**). Images acquired by ultrasound were analyzed using software resident on the machine. Ultrasound measurements taken were abdominal height, abdominal diameter, thoracic diameter, thoracic height, crown rump length, head length, and biparietal diameter (**Figure 1A**). Fetal sex was determined by ultrasonography on D77 of gestation (**Supplemental Figure 1**). Furthermore, fetal morphometry was assessed on D77 in a subset of 18 animals (6 AI and 12 ART animals). Ultrasound measurements taken on D77 were abdominal height, abdominal diameter, thoracic diameter, thoracic height, crown rump length, head length, and biparietal diameter.

### Surgical fetal collections of D56 and D105 fetuses

Heifers (n = 51 for D56; n = 48 for D105) were fasted at least 12 h prior to surgery. Fetuses were surgically retrieved to preserve nucleic acid integrity. All surgical procedures were performed by a licensed veterinarian.

### Collection of fetal tissues and fetal measurements of D56 and D105 fetuses

Collected fetuses and their fetal membranes were weighed, fetal morphometry was assessed, and abnormal phenotypes were noted. Measurements were crown-rump length, heart girth, forelimb length, biparietal length, abdominal height, head length and thoracic height. Subsequently, all tissues (i.e. pancreas, kidney, liver, ears, skeletal muscle, heart, diaphragm, tongue, buccal mucosa, umbiliculs, placenta [cotyledons and intercotyledon], brain, reproductive tract, gonad, skin, stomach, intestine, lung, spleen, tail, leftover carcass) were dissected and divided in two and immediately frozen in liquid nitrogen. For all collections, the same person measured and weighed all fetuses and fetal membranes, and another person (veterinary anatomic pathologist) dissected all the tissues. The average time from excision of the fetus from the uterus to when all tissues were frozen in liquid nitrogen was approximately 18 minutes. All tissues were stored at −86°C until further use.

### Image analysis of crown rump length and umbilicus diameter in D105 fetuses

Measurement of D105 fetuses’ crown rump length (from the top of the head to base of the tail) and diameter of the umbilicus (at the base of the umbilicus where it protrudes from the body) were measured using the ImageJ’s^42^ freehand line function using a lateral side picture of the D105 fetuses. The surface where the fetuses laid was squared (each square = 2.54 cm) and was used to convert all measurements to cm, and then a ratio of umbilicus diameter to crown rump length was determined.

### Maternal blood collection and processing

Maternal blood was collected via tail venipuncture on D55 and D105 of pregnancy into K3 EDTA tubes (BD, Franklin Lakes). Blood processing was done as described earlier in ^43^. Briefly, blood containing tubes were centrifuged at 1200 x *g* for 20 minutes at 4°C. The buffy coat was transferred to 15 milliliters centrifuge tubes containing 12 milliliters of red blood cell lysis buffer (150 mM NH_4_Cl, 10 mM NaHCO_3_, 1 mM EDTA, pH 7.0). White blood cells (WBC = leukocytes) containing tubes were briefly vortexed, incubated at room temperature for 5 minutes, and later centrifuged at 300 x *g* for 10 minutes at 4°C. After discarding the supernatant, the WBC pellet was washed once in 5 milliliters red blood cell lysis buffer and then with 5 milliliters ice-cold 1X Dulbecco’s phosphate buffered saline, with centrifugation at 300 x *g* at 4°C for 5 minutes at each wash. After discarding the supernatant, the WBC pellet containing tubes were placed immediately on dry ice and then stored at −86°C until use.

### Selection of samples for RNA sequencing (RNAseq)

The WBC samples of 23 D105 pregnant heifers were selected for RNAseq on the basis of their group (AI or ART), weight of ART fetuses (≥97 percentile of AI weight = ART-LOS or <97 percentile of AI weight = ART-normal), fetal sex, and whether the females carried one or two fetuses in the ART group. In addition, the D55 WBC samples of the same 23 females were also used for transcriptome analyses (Supplemental Figure 2).

**Figure 2.**
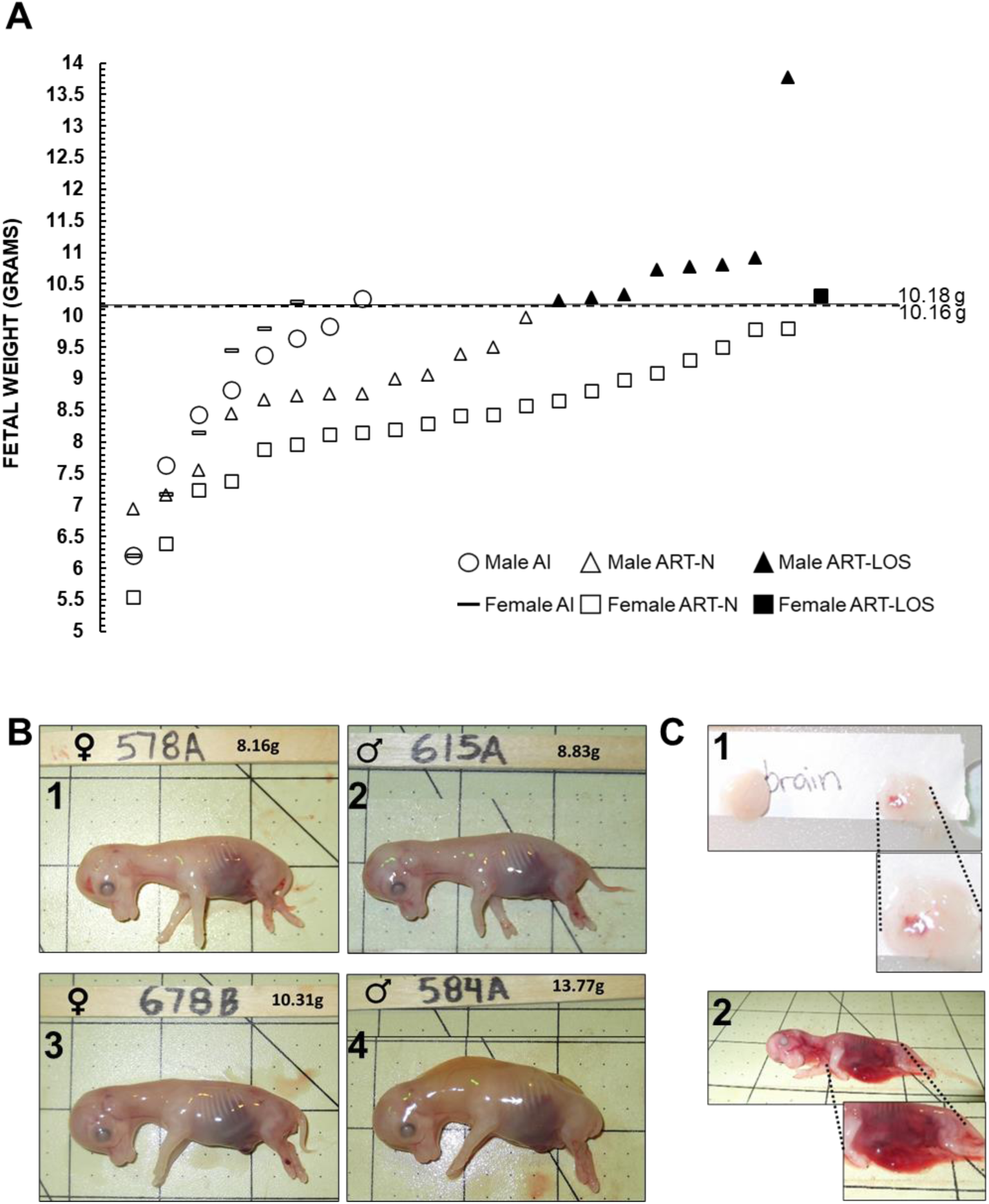
Day 56 fetal collections. (A) Fetal weight at D56 of gestation. The X axis has no actual implication and is used to scatter the spots representing each fetus for ease of visualization. The sex of the fetuses and the way they were generated is shown at the bottom of the figure. The bold line represents the 97 percentile of AI D56 fetal weight (i.e., male – 10.18 g, female − 10.16 g). (B) The pictures show D56 fetuses in the control (B1-B2) and ART-LOS group (B3-B4). Also, females (B1, B3) and males (B2, B4). B1 = AI-578A (control female weighing 8.16 g which is approximate average weight of female control fetuses). B2 = AI-615A (control male weighing 8.83 g which is the approximate average weight of male control fetuses). B3 and B4 show the heaviest LOS (ART-LOS 678B: female weighing 10.31 g and ART-LOS 584A: male weighing 13.77 g). Each square on the background = 2.54 cm. (C) Secondary phenotypes observed in day 56 LOS fetuses - focal hemorrhage on the brain (C1) and abdominal wall defects (C2). LOS: large offspring syndrome. AI: artificial insemination. ART: assisted reproductive technologies (i.e., *in vitro* produced embryos).

### RNA isolation and transcriptome analyses

Total RNA was isolated using Trizol™ reagent (Invitrogen, Carlsbad, CA) according to the manufacturer’s instructions and stored at −86°C until use.

#### Library preparation and transcriptome sequencing

RNA processing and sequencing was performed by BGI Americas Corporation (Cambridge, MA). Libraries were sequenced using the DNBSEQ-G400 platform to generate 20 million 100bp paired end reads.

#### RNAseq data analysis

Base quality and adapter contamination was assessed with FastQC and low quality (Pred < 20) raw reads trimmed using DynamicTrim. Trimmed reads less than 60 bases in length were removed with SolexaQA++ LengthSort function^44^. Retained paired-end reads were aligned to the bovine reference genome, ARS-UCD1.2^45^ with HISAT2 v2.1.0^46^ with the parameter adjustment (--mp 6,6; --score-min L,0,-0.2; --known-splicesite-infile) included to improve specificity. Total read counts for each gene were calculated by using the HTseq-count default union-counting module ^47^ using NCBI (GCF_002263795.1_ARS-UCD1.2) RefSeq gene set.

#### Hierarchical clustering

Raw read counts were normalized and the dist(method = ‘euclidean’) function was used to compute the distances between the rows of normalized count data. Finally, unsupervised hierarchical clustering was applied using the average linkage method and the base R hclust function to build the dendrogram.

#### Differential expression analysis

Differential expression analysis was performed using edgeR v3.32.1^48^ and DESeq2 v1.30.1^49^. Likelihood ratio tests were done using both packages (edgeR using glmLRT and DESeq2 using DESeq(test = LRT)). For edgeR, the raw read counts were normalized using RUVseq^50^. The betweenLaneNormalization function of EDAseq v2.24.0 was used to adjust for sequencing depth and then upper quantile normalization was done to remove unwanted variation^51^. The trimmed means of M values (TMM) was also used for edgeR analysis. For DESeq2, the median of ratios method of normalization was used using estimateSizeFactors() and assigning the normalization factors back to our count matrix. Pairwise comparisons between each treatment (Control-AI vs ART-Normal, ART-Normal vs ART-LOS, and Control-AI vs ART-LOS) group at each time point (D55 and D105) were performed. In addition, we compared the transcriptome of the dams carrying the two largest ART-LOS individuals (#604 and #664) against all other animals for D55 and D105. Furthermore, a pairwise comparison of WBC transcriptomes of females carrying two vs one fetuses was done to account for differential expression due to multiple fetuses (i.e. increased fetal mass). Genes with a false discovery rate (FDR; EdgeR) or adjusted P-value (padj; DEseq2) ≤ 0.05 were classified as significant. Significant genes from both packages were overlapped in order to generate a list of candidate genes for downstream analysis.

### cDNA synthesis and quantitative RT-PCR (qRT-PCR)

Total RNA was treated with DNase (Fischer Scientific, Waltham, MA) and used as template to synthesize cDNA in a 20 μl reaction using oligo dT and Superscript IV (Invitrogen, Carlsbad, CA) as recommended by the manufacturer.

*Arginine and Serine Rich Coiled-Coil 1* (RSRC1) and *Transcription Termination Factor 1* (TTF1) were used as test genes to corroborate RNA sequencing results based on their upregulation in the WBC transcriptome of the two females carrying the two largest LOS fetuses. Endogenous transcripts used as normalizers were: *Ecdysoneless Cell Cycle Regulator* (*ECD*), *Nuclear Factor of Kappa Light Polypeptide Gene Enhancer in B-Cells Inhibitor, Beta* (*NFKBIB*), and *VPS35 Endosomal Protein Sorting Factor Like* (*VPS35L*). These were chosen based on their constancy of expression in the RNAseq result (i.e., EdgeR FDR = 1; DESeq2 padj > 0.8 and coefficient of variation ≤ 0.10 across all 23 D105 WBC samples. In addition, these transcripts were also chosen based on availability of intron-spanning TaqMan probes (ABI, Foster City, CA) for bovine. TaqMan probe information is as follows: *ECD* (Bt03235022_m1), *NFKBIB* (Bt03247668_m1), *RSRC1* (Bt03236115_m1), *TTF1* (Bt03266651_m1), *VPS35L* (Bt03269522_m1). The mRNA levels of the target genes for the pregnant females carrying ART-LOS and the two largest LOS fetuses (cow #604 and #664) relative to the combined AI and ART-normal groups was calculated using the comparative cycle threshold (C_T_) method. Briefly, the C_T_ for each sample was normalized to the geometric mean of the three endogenous reference genes. The average C_T_ was calculated by averaging the C_T_ of all independent samples excluding those from the females carrying the two largest LOS fetuses. The comparative C_T_ method (ΔΔC_T_) was used to compare the values of 604 and 664 against the average C_T_ for all other samples. Fold difference is used for data representation.

### Statistical analyses

All analyses include an ART-normal group to remove potential confounding effects of method of conception when analyzing variables. The morphometric data were analyzed by analysis of variance using the general linear model procedure using SAS software v9.4 (SAS Institute, Cary, NC). Dependent variables were all fetal measurements, and the independent variables were the group and day of collection (AI, ART-normal, ART-LOS). Differences were considered statistically significance when p<0.10.

## RESULTS

### Pregnancy rate

The overall pregnancy rates on day 55 were 61.1% for the AI group and 43.0% for the ART group. For the D56 fetal collection set, there were 14 singleton pregnancies in the AI group, while, for the ART group, there were 19 singleton pregnancies and 12 females carried two conceptuses. For D105 fetal collection set, there were 12 singleton pregnancies in the AI group, while, for the ART group, there were 26 singleton pregnancies and 10 females carried two conceptuses.

### Day 55 fetal ultrasonographies

Fetal ultrasonographies were taken on gestation D55 to determine if LOS could be detected by this stage of pregnancy. Figure 1 shows the average measurements of all fetuses collected in this study (AI=26, ART=89). There were no differences in any of the fetal ultrasonographic measurements between the AI and ART groups. Head length was greater in male fetuses (n=69; mean ± SEM; p < 0.04; 2.01 ± 0.02) when compared to female fetuses (n=43; 1.94 ± 0.03). There was also a sex effect on crown rump length (p = 0.06) with males (4.64 ± 0.05) being longer than females (4.46 ± 0.06). This effect was more pronounced within the ART group (P < 0.03; 4.62 ± 0.05 and 4.42 ± 0.07, for males (n=53) and females (n=34), respectively).

### Fetal collections and LOS determination

Fetal overgrowth is the defining phenotype of LOS, and weight at surgical collection was used to define LOS using criteria described earlier by us^2^. A fetus was categorized as overgrown if its weight was ≥97 percentile weight of the AI in a sex-specific manner. We chose ≥97 percentile of control weight as threshold to ascribe LOS as this has previously been used to describe BWS^52^, the counterpart syndrome in humans.

### Day 56 fetuses

Fetal weight was not significantly different between singletons and twins (mean ± SEM; 8.94 ± 0.28 g and 8.79 ± 0.23 g, respectively). Fetal weight was similar between the AI and ART fetuses (mean ± S.D.; 8.66 ± 1.39 and 8.95 ± 1.43, respectively). For the AI group, there were eight males and six females. The average weight for the males was 8.78 ± 1.33 g (weight range: 6.21 - 10.27 g; Figure 2) while the average weight for the females was 8.50 ± 1.59 g (weight range: 6.20 - 10.22 g). For the ART group, we collected 21 males and 22 females. The average weight for the males was 9.52 ± 1.53 g (weight range = 6.94 - 13.77 g) while the average weight for the females was 8.40 ± 1.11 g (weight range = 5.55 - 10.31 g). Fetuses weighing ≥97 percentile of controls (male = 10.18 g and female = 10.16 g) in the ART group were considered LOS (Figure 2). In total, there were eight ART-LOS males (weight range = 10.23 - 13.77 g) and one ART-LOS female (weight = 10.31 g). All ART fetuses weighing <97 percentile weight of controls were referred to “ART-normal” (males – n=13 [weight range: 6.94 – 9.98 g] females – n=21 [weight range: 5.55 - 9.8 g]). Besides heavier body weight, other phenotypes observed in the ART group were focal hemorrhage on the brain and abdominal wall defects (Figure 2C). The fetal measurements at collection for the AI, ART-normal and ART-LOS groups are summarized in Figure 3. Only one female in the ART group was considered LOS, therefore no comparisons were made between it and the other groups.

**Figure 3.**
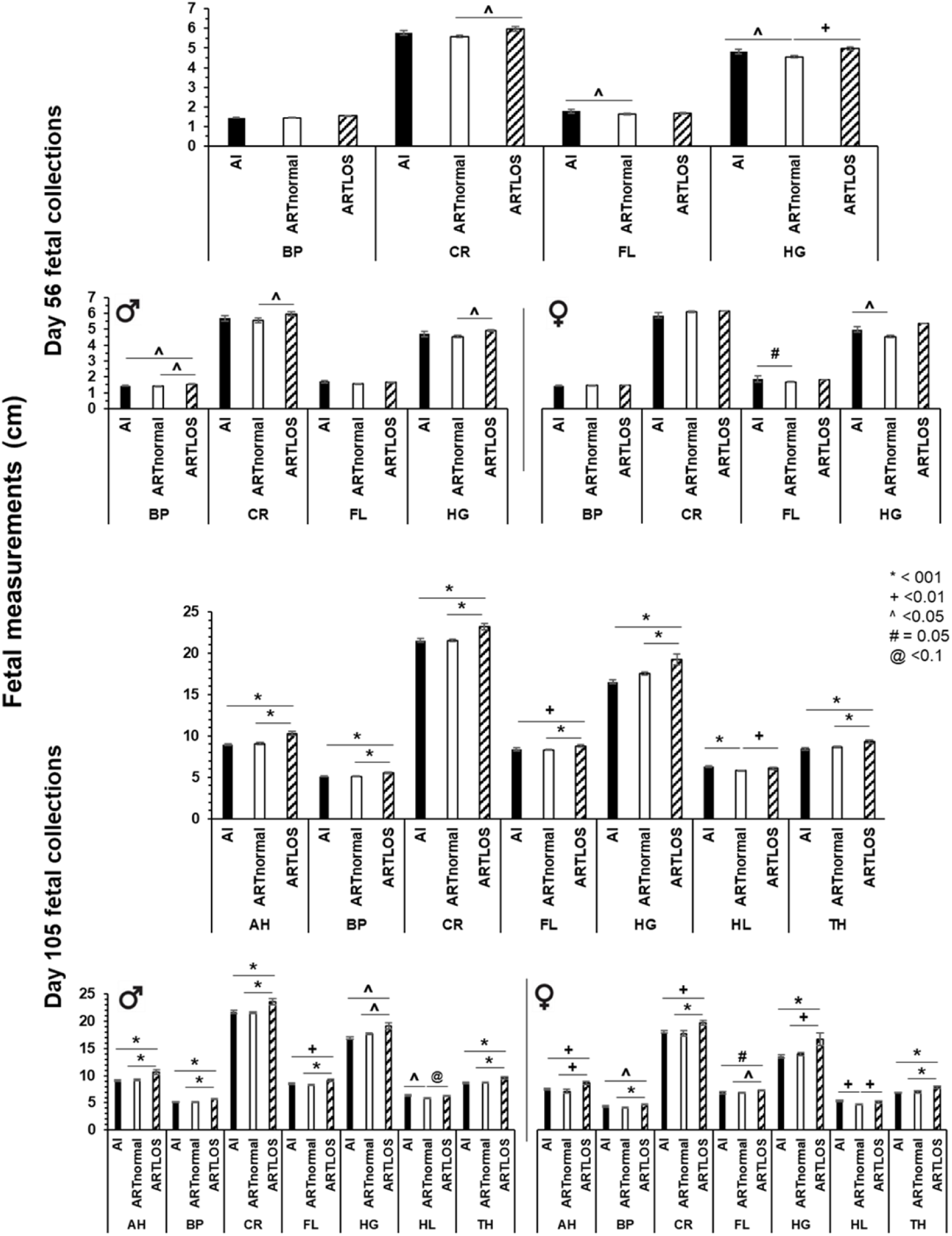
Fetal measurements at collection. Top three panels – D56 fetal measurements at collection (n= 6 females and 8 males in the AI group and 22 females [1 LOS] and 21 males [8 LOS] in the ART group). Bottom three panels – D105 fetal measurements at collection (n= 4 females and 8 males in the AI group and 13 [9 LOS] females and 33 males [8 LOS] in the ART group). For each day set, the top graph includes both sexes and the bottom two graphs are separated by sex, males on the left and females on the right. BP: biparietal diameter. CR: crown-rump length. FL: forelimb length. HG: Hearth girth. AH: Abdominal Height. HL: head length. TH: thoracic height. Data are represented as average ± SEM. Lines going over three bars are used to represent statistical differences between the first and the third bar.

### Day 105 fetuses

Fetal weight was not significantly different between singletons and twins (mean ± SEM; 532.78 ± 20.02 g and 494.80 ± 29.69 g, respectively). Fetal weight was similar between the AI and ART fetuses (mean ± S.D.; 466.00 ± 55.62 and 532.15 ± 134.80 g, respectively). For the AI group, we collected eight males and four females. The average weight for the males was 494.29 ± 44.05 g (mean ± S.D.; weight range: 442 - 550 g; note: there is a missed observation in this group) while the average weight for the females was 416.50 ± 36.01 g (weight range: 388 - 468 g; Figure 4). For the ART group, we collected 33 males and 13 females. The average weight for the males was 526.39 ± 123.00 g (weight range = 366 - 1080 g) while the average weight for the females was 546.77 ± 167.77 g (weight range = 318 - 986 g). Fetuses weighing ≥97 percentile of controls (male = 548.92 g and female = 463.14 g) in the ART group were considered LOS (Figure 4).

**Figure 4.**
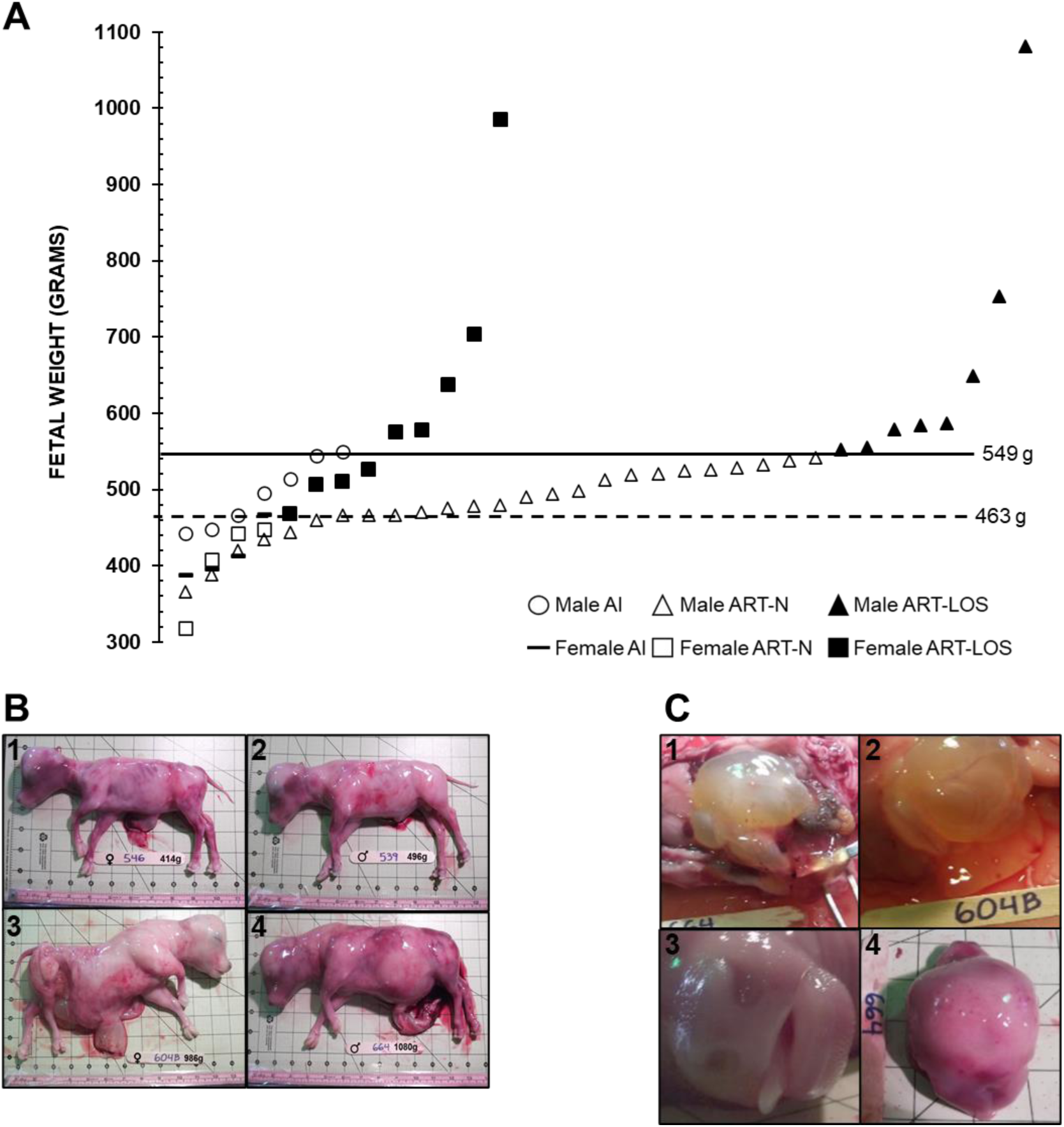
Day 105 fetal collections. **(A)** Fetal weight at D105 of gestation. The X axis has no actual implication and is used to scatter the spots representing each fetus for ease of visualization. The sex of the fetuses and the way they were generated is shown at the bottom of the figure. The bold line represents the 97 percentile of AI D105 fetal weight (i.e., male – 549 g, female - 463 g). **(B)** The pictures show D105 fetuses in the control (B1-B2) and ART-LOS group (B3-B4). Also, females (B1, B3) and males (B2, B4). B1 = AI-546 (control female weighing 414 grams which is approximate average weight of female control fetuses). B2 = AI-539 (control male weighing 496 grams which is the approximate average weight of male control fetuses). B3 and B4 show the heaviest LOS (ART-LOS 604B: female weighing 986 grams and ART-LOS 664: male weighing 1080 g). Each square on the background = 2.54 cm. **(C)** Secondary phenotypes observed in day 105 LOS fetuses - gelatinous material in the peritoneal cavity and organs (C1-2) long protruding tongue (C3), skull asymmetry (C4). AI: artificial insemination. ART: assisted reproductive technologies (*i*.*e*., *in vitro* produced embryos).

In this study, it appears that female fetuses were preferentially lost between D56 and D105, especially in the ART group. On day 56, we collected 6 females and 8 males in the AI group and 22 females and 21 males in the ART group. On D105 we collected 4 females and 8 males in the AI group and 13 females and 33 males in the ART group, which is different than the expected 1:1 ratio (p < 0.01). Although the reason for the loss of female fetuses between D56 and D105 is unknown, this was far more pronounced in the ART group. The deficiency of female fetuses was not statistically significant in the AI group (p = 0.20) but was highly significant in the ART group (p < 0.003) when compared to the binomial distribution with the expected 1:1 sex ratio.

In total, there were eight ART-LOS males (weight range = 552 - 1080 g) and nine ART-LOS females (weight range = 468 - 986 g). All ART fetuses weighing <97 percentile weight of controls were referred to “ART-normal” (males – n=25 [weight range: 366 – 542 g] females – n=4 [weight range: 318 - 448 g). Besides heavier body weight, other phenotypes observed in the ART group were long protruding tongue (Figure 4C.3), large organs (heart, kidney, lung, pancreas), hepatic cyst, abdominal ascites, gelatinous material in the peritoneal cavity and organs (Figure 4C.1-2) and skull asymmetry (Figure 4C.4). In addition, large umbilicus (Figure 4B.3) were observed at collection in two of the female ART-LOS fetuses (fetus number 656 and 604B), however, the ratio of the umbilicus to the crown-rump length (as determined by image analysis) was similar between groups, although the largest female (604B) had at least a 40% wider base of the umbilicus when compared to all other fetuses (Figure 4B.3). There was a group (i.e. AI, ART-normal, ART-LOS) effect for crown-rump length, abdominal height, biparietal diameter, forelimb length, heart girth, thoracic height and head length (p < 0.002) and sex effects for crown-rump length, abdominal height, biparietal diameter, forelimb length, and thoracic height (p < 0.05). The fetal measurements at collections are summarized in Figure 3.

### Associations between D55 fetal ultrasonographies and collection measurements

#### D55 ultrasonographic measurements on fetuses collected on D56

The summary of D55 ultrasonographic measurements for fetuses collected on D56 may be found in Figure 5. Female specific analysis for the ART-LOS group was not performed as only one female in the ART group weighed more than the ≥97 percentile threshold used to ascribe overgrowth. No differences were observed for sex, group, or their interaction for abdominal diameter, abdominal height, biparietal diameter, thoracic diameter, or crown-rump length. Head length was greater in males than females (p < 0.006; Mean ± SEM; 2.05 ± 0.02 and 1.93 ± 0.03 for males and females, respectively). A group by sex interaction was detected for thoracic height in which the males of the ART-normal group were smaller than males in the AI and ART-LOS groups (p < 0.03; Mean ± SEM; 1.60 ± 0.04, 1.75 ± 0.05 and 1.83 ± 0.08 for ART-normal, AI and ART-LOS groups, respectively).

**Figure 5.**
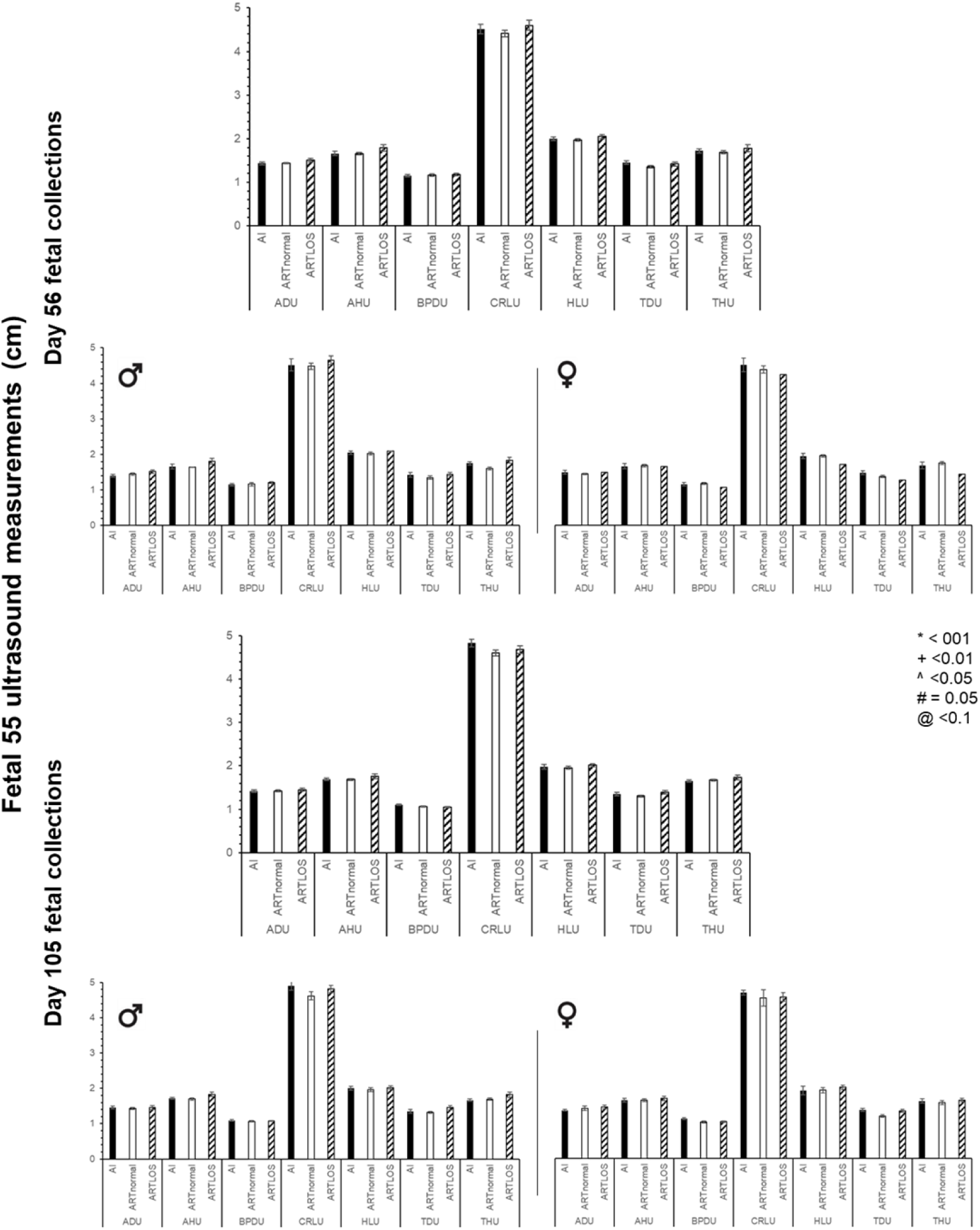
Group-specific day 55 fetal ultrasound measurements. Top three panels – D55 ultrasonographic measurements of fetuses collected on D56. n= 6 females and 8 males in the AI group and 22 females [1 LOS] and 21 males [8 LOS] in the ART group. Bottom three panels – D55 ultrasonographic measurements of fetuses collected on D105. For each day set, the top graph includes both sexes and the bottom two graphs are separated by sex, males on the left and females on the right. n= 4 females and 8 males in the AI group and 13 [9 LOS] females and 33 males [8 LOS] in the ART group. U: ultrasound. AH: abdominal height. AD: abdominal diameter. BPD: biparietal diameter. CRL: crown rump length. HL: head length. TD: thoracic diameter. TH: thoracic height. Data are represented as average ± SEM. For D56, there was only one female considered LOS, hence the lack of error bars. Lines going over three bars are used to represent statistical differences between the first and the third bar.

A slight positive correlation was observed between the day 55 ultrasonographic measurements and the D56 fetal weight for abdominal diameter (0.40; p < 0.003), abdominal height (0.36; p < 0.007), crown-rump length (0.34; p < 0.01), head length (0.32; p < 0.02), and thoracic height (0.25; p = 0.06), while no correlations were observed between biparietal diameter or thoracic diameter measurements. When males and females were analyzed separately, positive correlations were observed for abdominal diameter (0.57; p < 0.002), abdominal height (0.64; p < 0.0003), crown-rump length (0.36; p = 0.06), and thoracic height (0.54; p < 0.004) in males. However, no correlations were found between any of the D55 ultrasonographic measurements and D56 fetal weight in females.

#### D56 ultrasonographic measurements on fetuses collected on D105

The summary of D55 ultrasonographic measurements for fetuses collected on D105 may be found in Figure 5. No differences were observed for sex, group, or their interaction for abdominal diameter and head length. Females were smaller than males in their crown-rump length (p = 0.08), abdominal height (p < 0.04), thoracic diameter (p = 0.07), and thoracic height (p < 0.008).

A moderate positive correlation was observed between the D55 ultrasonographic measurements and the D105 fetal weight for abdominal diameter (0.57; p < 0.0001) and abdominal height (0.58; p < 0.0001). A slight positive correlation was observed between fetal weight and crown-rump length (0.27; p <0.04), head length (0.33; p < 0.02), thoracic diameter (0.34; p < 0.02), and thoracic height (0.49; p < 0.0002) while no correlation was observed for biparietal diameter. For males, there was a moderate positive correlation between fetal weight and abdominal diameter (0.52; p < 0.0009), abdominal height (0.57; p <0.0002), and thoracic height (0.56; p < 0.0003) and slight positive correlation for thoracic diameter (0.36; p < 0.03). For females, there was a moderate positive correlation between fetal weight and abdominal diameter (0.67; p < 0.005), abdominal height (0.67; p < 0.005), and head length (0.59; p < 0.02).

#### D77 ultrasonographic measurements on fetuses collected D105

An attempt was made to determine fetal morphometry on the subset of day 77 pregnant group (AI= 6; ART=12), however this was not possible or reliable for many of the samples as the fetus was too large to do accurate measurements (data not shown).

#### Are D55 ultrasonographic measurements useful to identify LOS?

Overall, no single ultrasonographic measurement can explain LOS on day 105 of gestation. We tested various combinations of measurements and identified a strong positive correlation between D105 fetal weight and the product of the D55 ultrasonographic measurements for abdominal diameter, abdominal height, crown-rump length, head length, thoracic height, and thoracic diameter (0.76; p < 0.007 and 0.72; p < 0.0001 for AI and ART fetuses, respectively; Figure 6). The highest number resulting from the multiplication of the beforementioned ultrasonographic measurements was 79.92. The D105 ART fetuses were compared to that threshold and all except the two most extreme LOS cases (fetus 604B and 664 weighing 986 and 1080 g, respectively) were on or below the AI threshold. Similar comparisons were made for the set of fetuses collected on D56. While the correlation was also strong (0.75; p < 0.003), the threshold for the AI was higher (91.32) than for that obtained for the D105 AI fetuses. The correlation decreased to 0.46 (p < 0.03) for the ART group indicating more variability in fetal weight at this stage and perhaps inclusion of fetuses that will be lost later during pregnancy or will have a differential rate of growth after this stage. In this group, only the heaviest ART fetus was above the 100-threshold used in the D105 group.

**Figure 6.**
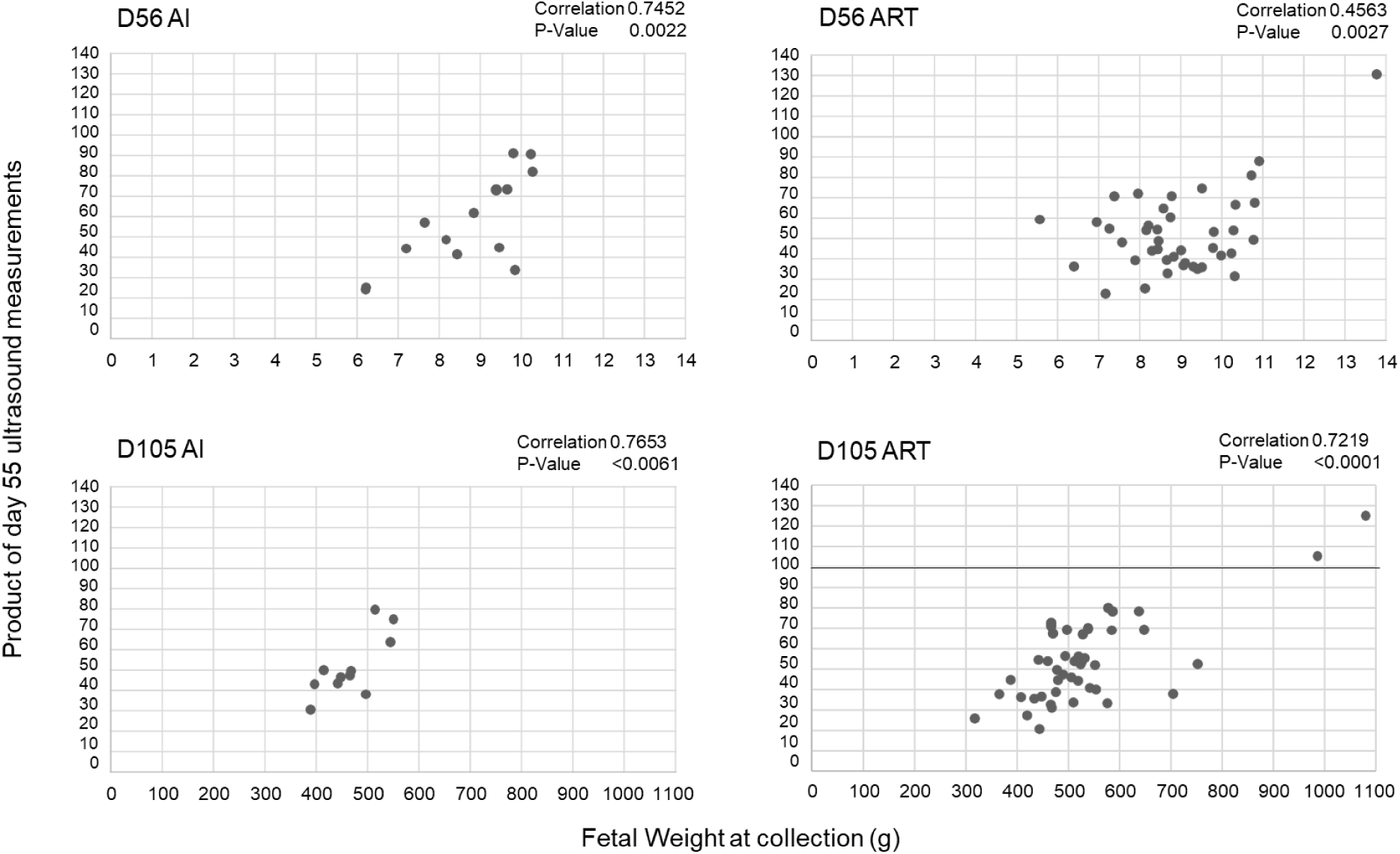
Correlation between fetal weight at collection and the product of D55 ultrasonographic measurements. **(A)** D56 AI group. **(B)** D56 ART group. **(C)** D105 AI group. **(D)** D105 ART group. Ultrasonographic measurements included = abdominal diameter, abdominal height, crown-rump length, head length, thoracic height, and thoracic diameter. AI: artificial insemination. ART: assisted reproductive technologies (*i*.*e. in vitro* produced embryos).

#### Maternal blood transcriptome analysis

Only reads which aligned to known genes of the bovine reference genome assembly ARS-UCD1.2 using NCBI (GCF_002263795.1_ARS-UCD1.2) were used in the present study (Supplementary Table 1).

Unsupervised hierarchical clustering of the normalized read counts showed 18 of 46 samples (ie. D55 and D105 WBC transcriptomes of the same 23 females) clustered by animal (Supplementary Figure 3). In other words, the D55 and D105 samples from 9 of the heifers grouped together by individual and this was irrespective of treatment group (AI, ART-normal or ART-LOS). In addition, the analysis separated the females carrying the two largest D105 ART-LOS fetuses (dam #604 and #664) from the rest of the animals (Supplementary Figure 3). The day-specific unsupervised hierarchical clustering analyses may be found in Supplementary figure 4.

Figure 7 shows the results for differentially expressed genes identified during the pairwise comparisons between each treatment group at each time point (D55 and D105). In addition, a pairwise comparison of WBC transcriptomes of females carrying two fetuses vs one was done to account for differential expression due to multiple fetuses (i.e. increased fetal mass). Furthermore, we compared the transcriptome of the dams carrying the two largest ART-LOS individuals (#604 and #664) against all other animals for D55 and D105. Overall, for the D55 comparison, there were 13 differentially expressed genes identified by EdgeR and 8 identified by DESeq2 and for D105 there were 31 differentially expressed genes identified by EdgeR and 4,451 by DESeq2. Data show that a large number of genes identified as differentially expressed are uncharacterized transcripts (“LOC”). Maternal WBC transcriptome analyses found that *LOC783838* and *PCDH1* were identified as differentially expressed in the extreme cases of LOS on gestation days 55 and 105 by both EdgeR and DESeq2 statistical packages (Figure 7). In addition, transcript levels of *ACTA2, KDM5A, MAN1A2, MIR2376, PRRC2C, RSBN1, S100A14, SRPK2*, and *TTF1* were identified as differentially expressed in the females carrying the two largest fetuses on D105. For qRT-PCR corroborations, we focused on two genes upregulated in the two largest LOS fetuses when compared to all other fetuses, and whose intron-spanning TaqMan probes were readily available, namely *TTF1* and *RSRC1. TTF1* was identified as differentially expressed by both EdgeR and DESeq2 statistical packages and *RSRC1* was identified as differentially expressed by DESeq2. Data show that the pattern of expression for these genes is similar between RNAseq and qRT-PCR results (Figure 8).

**Figure 7.**
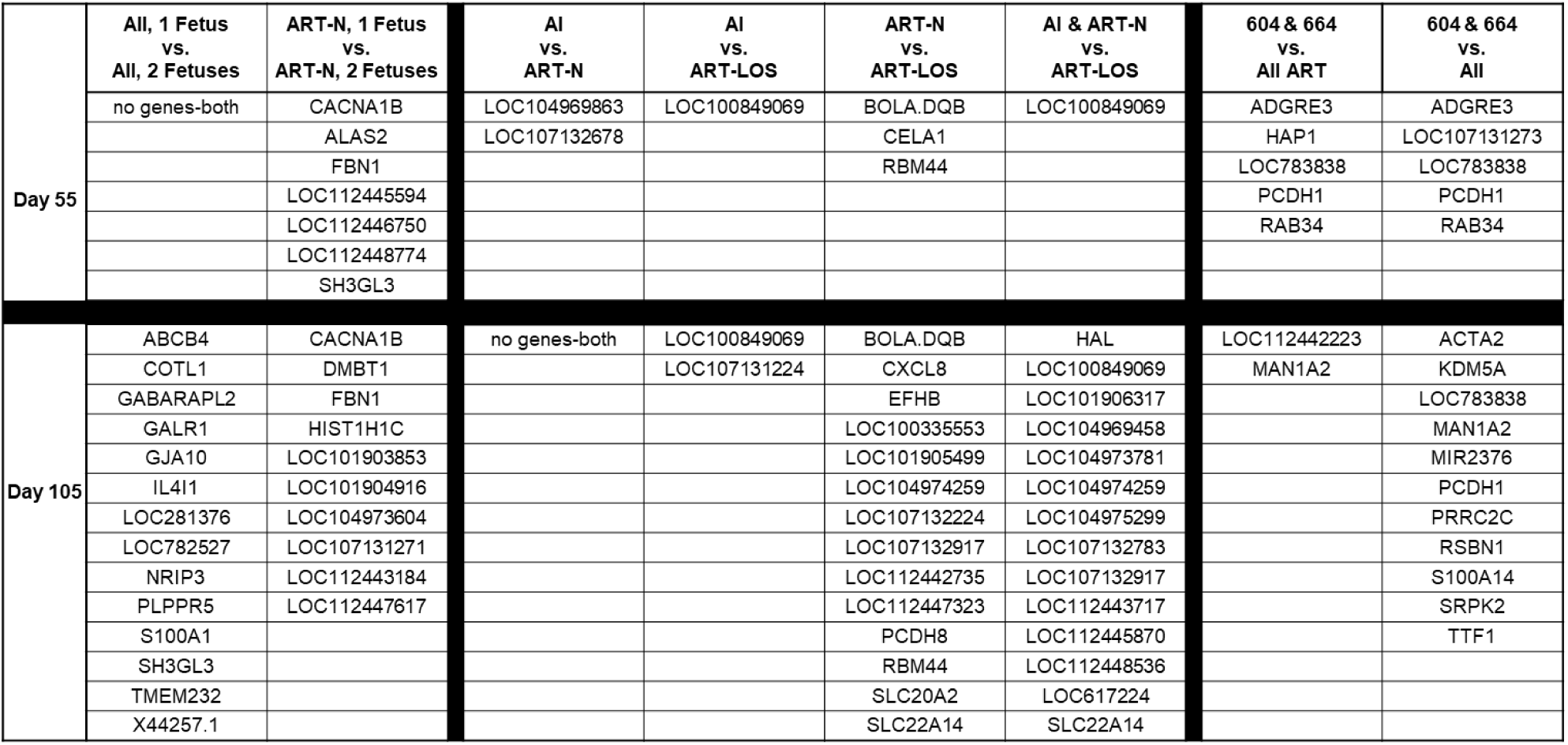
Differentially expressed genes in D55 and D105 maternal WBC transcriptomes. Shown are the genes identified by both EdgeR and DESeq2 as statistically different (p < 0.05) in the named comparisons. AI = artificial insemination (i.e. control). ART-N = embryos were produced by *in vitro* procedures that were <97% of the control’s weight at D105. ART-LOS embryos were produced by *in vitro* procedures and were ≤97% of the control’s weight at D105. “604 & 664” are the numbers of the heifers carrying the two largest LOS fetuses. Gene names starting with “LOC” are uncharacterized transcripts. The “No gene-both” designation notes that genes were identified as differentially expressed by either EdgeR or DESeq2 but not both.

**Figure 8.**
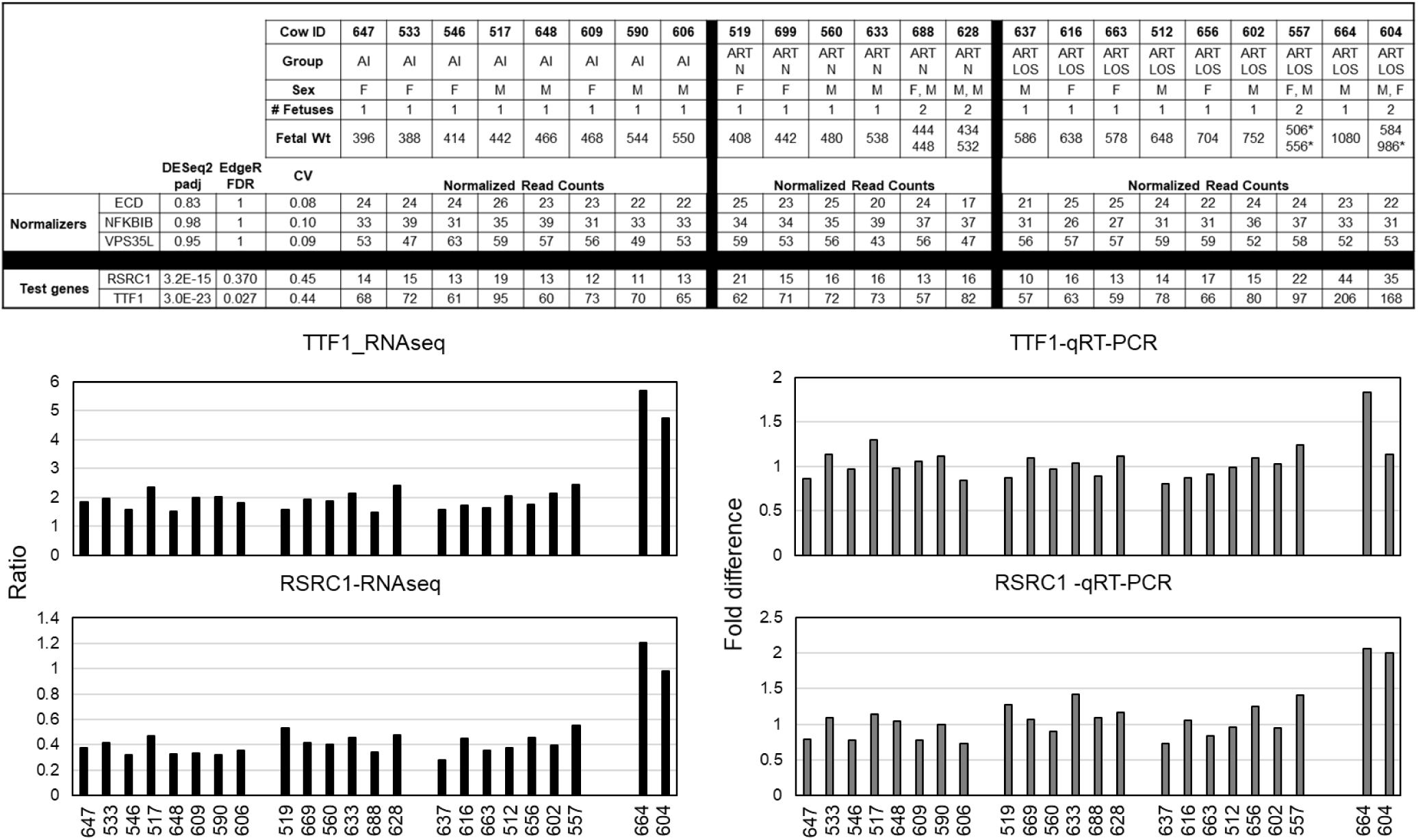
Normalized read counts and qRT-PCR corroborations of D105 maternal transcriptomes. **Top**. Table showing all information pertaining the samples and the transcriptome’s statistics for the genes chosen for qRT-PCR corroborations. The table is organized by total fetal mass in increasing order. # of fetuses indicates how many fetuses the dam was carrying at the time of collection. F = female fetus. M = male fetus. AI = artificial insemination (*i*.*e*. control). ART-N = embryos were produced by *in vitro* procedures that were <97% of the control’s weight at D105. ART-LOS embryos were produced by *in vitro* procedures and were ≤97% of the control’s weight at D105. CV = coefficient of variation of all 23 samples. Asterisk – denotes the LOS of the pair. **Bottom**. Bar graphs showing RNAseq results represented as sample ratio of the named gene from the mean of the three normalizers and qRT-PCR results represented as fold difference. For qRT-PCR, the geometric mean of three endogenous transcripts, namely *ECD, NFKBIB* and *VPS35L* were used to normalize the levels of *TTF1* and *RSRC1*. Data are represented as fold difference from the average of 21 samples (average fold difference = 1; not including 604 and 664). Transcripts of *TTF1* and RSRC1 for 604 and 664 were compared to the normalized average of 21 samples for each test gene (ΔΔC_T_).

## DISCUSSION

The goal of the present study was to determine the usefulness of D55 of gestation fetal morphometry and D55 and D105 maternal leukocyte transcriptome for the identification of congenital fetal overgrowth in cattle. In total, 20.93% (9/43) of the D56 and 36.95% (17/46) of the D105 collected ART fetuses were considered LOS (≥97 percentile of the sex-specific weight of controls). The study revealed that the product of the D55 ultrasonographic measurements for abdominal diameter, abdominal height, crown-rump length, head length, thoracic height, and thoracic diameter may be used as an indicator of extreme cases of LOS. In addition, we identified that LOS fetuses had a different growth pattern from AI and ART-normal fetuses after D55 of gestation and that LOS fetuses had several developmental abnormalities such as hemihyperplasia (asymmetric growth), enlarged tongue, brain hemorrhage, enlarged umbilical cord, abdominal ascites, and abdominal wall defect. Maternal leukocyte transcriptome analyses identified that *ADGRE3, LOC107131273, LOC783838, PCDH1*, and *RAB34*, were differentially expressed on D55 in the females carrying the two largest LOS fetuses. On D105 of pregnancy the leukocyte transcriptomes of the same females had differential expression of *ACTA2, KDM5A, LOC783838, MAN1A2, MIR2376, PCDH1, PRRC2C, RSBN1, S100A14, SRPK2*, and *TTF1* when compared to all other animals.

Collection of fetuses was done on D56 of gestation since organogenesis has been shown to be completed before this day in cattle^53^. The reason behind this decision was to answer other aspects of the project, which are beyond the scope of the current study, such as questions regarding the epigenetic mechanisms associated with abnormal organ formation in LOS; information that will be used to identify etiologies of fetal overgrowth syndrome in cattle and human (i.e. BWS). Furthermore, fetuses were also collected on D105 as we have previously shown that fetal overgrowth is evident at this stage of gestation^2^. In humans, 97 percentile criteria is used to describe macrosomia in newborn babies^52^. Since macrosomia is the main characteristic of LOS and one characteristic that can result in dystocia, we used this weight to ascribe LOS in the present study, similar to what we have done previously.

A greater than two-times increase in fetal weight was observed in two of the D105 ART-LOS fetuses in the current study, an observation also reported by others^4^, suggesting that if those fetuses were allowed to go to term, dystocia would be probable and assisted delivery or caesarean section would be required. Dystocia as a result of LOS can lead to neonatal death^4^ and/or death of cows^22^. Further, dystocia associated with stillbirth^54^, is a major contributor to perinatal mortality in cattle^55^, and has also been shown to increase the chances of metritis^56^, and lameness^28^. Furthermore, dystocia can have an adverse impact on milk production^57^ and increase the calving to conception interval in dairy cows^58^. Thus, overgrown fetuses can have negative economic consequences to cattle producers^22^ and identification of those large calves during early pregnancy would help overcome these problems.

In the current study, we performed fetal ultrasonographic measurements at D55 for fetuses that were collected on D56 or D105 of gestation. Based on our findings, it is evident that the ≥97 percentile AI weight criteria used to assign a fetus as being LOS while useful at D105, it is not appropriate at gestation D56. From the findings of the correlation analysis between fetal weight and the product of six ultrasound measurements, we hypothesize that fetuses are lost after this day of pregnancy and/or will have subsequent disproportionate growth. This is in accordance to previous work which showed that smaller *in vitro* or cloned fetuses in the first trimester resulted in heavier fetuses at term^7,59^. Further, Bertolini and coinvestigators also suggested that *in vitro* produced fetuses show early growth retardation and then follow acceleration in fetal growth at later stages of gestation, showing biphasic growth pattern^7^.

Previous research reported that larger biparietal diameter might be a useful measurement to identify LOS at day 63 of gestation in cloned LOS fetuses^60^ and that a smaller crown rump length might be a useful measurement to identify LOS at day 58 of pregnancy in *in vitro* produced LOS fetuses^7^. However, those measurements were not found to be indicators for LOS in our current study. Possibilities for the discrepancies in our findings, are; 1) different definition of LOS (we used ≥97% weight of the control fetuses while Bertolini used fetuses that were 33% heavier than controls for comparisons^7^, 2) improvement in ultrasonography technology allowing for more accurate measurements, and 3) number of observations (113 fetuses in our study vs. 34 fetuses in^59^).

When we tried to do fetometry at day 77 of gestation, we were only able to measure head length with some accuracy. All other measurements were not reliable, indicating that day 77 ultrasonography might be too late to try to accurately identify LOS. Given the allometric growth that occurs after D56 and the fact that day 77 is too late to predict LOS by ultrasonography, and that fetal sex is most accurately predicted between days 60-80^34^, we suggest that future studies focus on day ∼65 as a target day to identify LOS in a sex-specific manner.

Larger than normal umbilicus and presence of large amounts of fluid-gelatinous material in the abdominal cavity were observed in largest/heaviest D105 ART-LOS fetuses. This is similar to what Constant *et al*.^59^ reported in day 220 fetuses produced by somatic cell nuclear transfer. In that study, the authors suggested that a large umbilical cord and abdominal ascites were not the result of fetal overgrowth per se, but rather a consequence of placental dysfunction, which in turn led to placental overgrowth. Contrary to this, other studies have shown an association of placental defects in cloned fetuses, with their loss during early pregnancy as a result of growth retardation^61,62^. Taken together, these studies suggest that fetuses with severe placental defects may be lost during early pregnancy and if those fetuses with placental defects survive, they could have higher placental and fetal growth at later stages of pregnancy through compensatory mechanisms. In the current study, the conceptuses were surgically removed to allow rapid collection of tissues in order to preserve nucleic acid integrity for other aspects of the project, therefore, even though we collected the placentas, we were not able to make thorough morphological assessments of this tissue. Regardless, no obvious placental abnormalities were evident at collection for ART conceptuses.

Enlarged tongue was also observed in D105 ART-LOS which is similar to previous findings in our laboratory^2,23^ and comparable to what has been observed in a similar congenital overgrowth condition in humans, namely Beckwith-Wiedemann Syndrome^63^. Large tongues can lead to difficulty in suckling and increase the chances of prenatal death^26^. In addition, the largest D105 ART-LOS fetus showed brachycephaly and asymmetrical growth of the cranium, an interesting finding given that one characteristic of BWS is hemihyperplasia^63^. These similarities demonstrates that BWS and LOS are the same syndrome, as previously reported by us^2^, and that they share similar misregulated developmental epi(genetic) mechanisms associated with asymmetrical growth.

In our study, we also had the objective of determining if maternal blood could be used as a biomarker to identify LOS on D55 and/or D105 of pregnancy. For this, we analyzed leukocyte transcriptome of 23 females carrying D105 AI, ART-normal (weight) and ART-LOS fetuses. We also analyzed the D55 leukocyte transcriptomes of the same females. Our initial approach was to do an unsupervised hierarchical clustering of de-identified samples to determine if obvious difference existed between the females carrying LOS fetuses when compared to the other two groups. To our surprise, the transcriptome of 18/46 samples clustered together by animal (i.e., n=9) regardless of pregnancy stage (i.e., D55 and D105). Another interesting point is that the experiment was done over three seasons (Autumn, Spring and Summer) with a range in temperature of −30°C to 43°C, and this was not detected in the transcriptome given the clustering by individual. Further, given the design of the study, in which we transferred two embryos per recipient heifer, some pregnancies in the D55 and D105 ART groups had two fetuses, however, the unsupervised hierarchical clustering did not cluster animals by number of fetuses. The unsupervised hierarchical clustering, did however, cluster the D105 transcriptomes of the females carrying the two largest LOS from all other females.

For the maternal leukocyte transcriptome analysis, we focused on the two females that carried the largest LOS fetuses as those were the ones that separated by hierarchical clustering and the ones that would most likely cause a difficult birth. Analyses identified *ADGRE3, LOC107131273, LOC783838, PCDH1*, and *RAB34*, as being different on D55 and *ACTA2, KDM5A, LOC783838, MAN1A2, MIR2376, PCDH1, PRRC2C, RSBN1, S100A14, SRPK2*, and *TTF1* on D105 when compared to all other animals. For qRT-PCR corroborations we focused on two transcripts whose TaqMan probes were readily available, namely *TTF1* which was identified as differentially expressed on D105 by EdgeR and DESeq2 statistical packages as well as *RSRC1* which was identified as differentially expressed by DESeq2. Analyses show consistency of expression between RNAseq and qRT-PCR results. Other genes will be tested in the future as assays become available.

Our study has several limitations. As it pertains to fetal ultrasonographies, published work shows that fetal growth varies among different breeds^64^ and that size difference can be noticeable as early as 3 months of gestation, therefore it is possible that fetal growth patterns may vary as early as D55 of gestation among different breeds; however, there is lack of published data showing that on D55 of gestation, fetuses from different breeds differ in size. Future research would have to address this question. Here, we also did sex-specific analyses for all our variables, including for ultrasonographic measurements. Since in our experiment, the number of male fetuses was unexpectedly higher in the ART group at D105, female specific ultrasonographic data analysis may be limiting. Finally, and importantly, it should be noted that during transcriptome analysis, we only used the known (mostly coding) group of bovine transcripts for this study and that the non-coding and novel transcript portion of the transcriptome remains unexplored. Future work will focus on these types of transcripts.

In summary, here we document initial efforts to identify LOS during the first trimester of pregnancy in cattle. We found that the product of the D55 ultrasonographic measurements for abdominal diameter, abdominal height, crown-rump length, head length, thoracic height, and thoracic diameter may be useful to identify the largest fetuses, whereas maternal leukocyte transcriptome analyses suggest *LOC783838* and *PCDH1* as potential markers for extreme cases of LOS on gestation days D55 and D105. In addition, transcript levels of *ACTA2, KDM5A, MAN1A2, MIR2376, PRRC2C, RSBN1, S100A14, SRPK2*, and *TTF1* may also serve as biomarkers on D105 of pregnancy for extreme cases. In addition, our analysis identified several genes that were misregulated in all LOS fetuses when compared to fetuses of normal weight produced by ART, future work will query the usefulness of these genes to predict milder cases of LOS. Finally, the long-term goal of this research is to identify the best time of pregnancy to capture LOS by ultrasonography and train a model using fetuses that will be allowed to go to term to determine the best maternal blood markers to identify fetal overgrowth in cattle.

## Acknowledgements

We would like to thank Drs. Martha Sofia Ortega, Dave Patterson, Jordan Thomas, and Mike Smith for their help during estrous synchronization and technical assistance through the project and Kenneth Ladyman and Reinhard Van Zyl for farm management and related help. We also thank other members of the Rivera laboratory (Ali Patten, Amanda Moreno, Monique Ferrell, Casper Safranski, Olivia Styron, and Carla Reyes) for help with preparations and tissue collections. We would also like to thank Cooper Stansberry and Michael Campbell for their help during oocyte collection and Dr. Ky Pohler for facilitating the donation of the Brahman semen used in the project. We would also like to thank Drs. Cliff Miller and Dr. Neal Martin’s veterinary technicians Sarah K., Carolyn W. and Chase H for veterinary assistance. This project is supported by the Agriculture and Food Research Initiative competitive grants no. 2018-67015-27598 and 2021-67016-33417 from the United Stated Department of Agriculture National Institute of Food and Agriculture and the L.E. “Red” Larson Endowment (PJH).

## Authors contributions

RMR designed the experiment. RMR, BNP, AKG, DEH, EJSM, YL, CK, CM, FWIII, EJ, YX, PT, EEC, ARBR, PJH, ZW, CMS, NM, CGE conducted the experiment and collected data. RMR wrote the original draft and all authors contributed to the manuscript.

## Data Availability Statement

The raw FASTQ files are publicly available at Gene Expression Omnibus (GEO accession no. GSE179946).

## Supplemental Figures

**Supplemental Figure 1.**
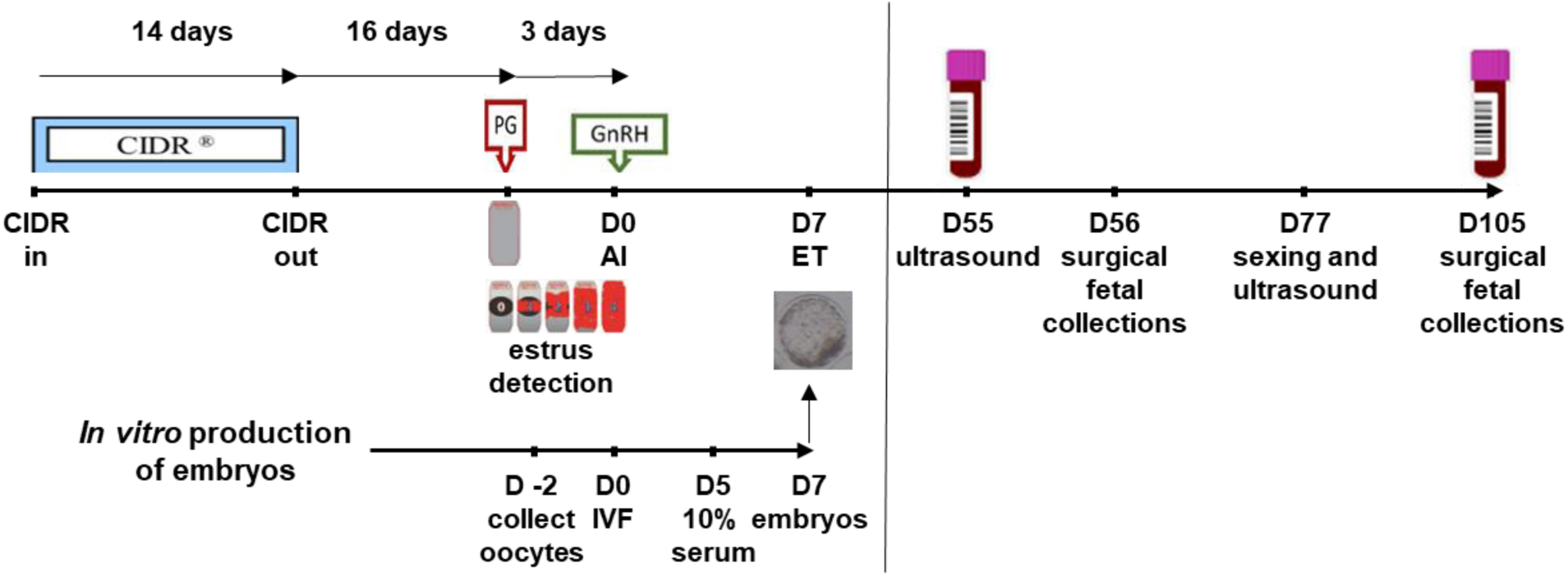
Experimental design. Production of day 56 and day 105 fetuses. For estrus synchronization, the 14-day CIDR®-PG & TAI protocol was followed. Briefly, CIDRs were placed in the heifers and removed 14 days later. Sixteen days after CIDR removal, 25 milligram of prostaglandin F2 alpha (Lutalyze); Zoetis, NJ) was administered intramuscularly. Concurrent with the administration of prostaglandin F2 alpha, a breeding indicator patch (Estrotect, Genex, Shawano, WI) was applied to each animal across the backbone as per the manufacturer’s instructions. Estrus was checked three times per day (7:00 h, 12:00 h and 16:00 h/17:00 h) for three consecutive days and only animals with a heat score two and above at the time of artificial insemination (AI; 13:00 h) were selected for AI or embryo transfer (ET). The breeding indicators were scored 0-4, with a score of 0 indicating no patch activation; a score of 1 signifying <25% patch activation; a score of 2 signifying >25 to <50% patch activation; a score of 3 signifying >50 to <75% patch activation; and a score of 4 signifying >75% patch activation. Heifers were randomly assigned to the AI or ET group. Regardless of experimental group, all animals were injected with 100 microgram of gonadotropin releasing hormone (GnRH) intramuscularly (Factrel, Zoetis) at the time of corresponding to the insemination in the AI group. In order to collect the number of fetuses required for the experiment, two sets of estrus synchronizations were performed (in November of 2018 and in February of 2019). *In vitro* production (IVP) of bovine embryos was done simultaneously with estrus synchronization to ensure AI (control) and IVP embryos were of the same chronological age on day 7 after estrus. Media and procedures were as previously described by us. Briefly, Bos *taurus taurus* (*B. t. taurus*; Angus/Angus-Crossbred) ovaries were obtained at an abattoir and oocytes collected at Oklahoma State University (OSU) in Stillwater Oklahoma. Oocytes were placed in CO2 equilibrated maturation medium and shipped overnight at 38.5°C to the University of Missouri (MU) or the University of Florida (UF). In addition, *B. t. taurus* oocytes were also purchased from DeSoto Biosciences (Seymour, TN, USA) and processed at MU. All media for embryo production were prepared at MU by a single technician and shipped overnight to the pertinent location prior to each procedure. Two sources of oocytes and two *in vitro* production (IVP) locations were used to ensure sufficient embryos were available for embryo transfer in case of technical or weather-related difficulties. Oocytes were removed from maturation medium after ∼21 h of culture and inseminated with semen from one *B. t. indicus* male (Brahman breed [JDH MR MANSO 7 960958 154BR599 11200 EBS/INC CSS 2]). Putative zygotes were stripped of cumulus cells by five minutes vigorous vortexing at approximately 18 h after insemination and cultured in KSOM supplemented with amino acids in a humidified atmosphere containing 5% O2, 5% CO2, and 90% N2. On day five after insemination, the culture medium was supplemented with 10% (v/v) estrus cow serum (collected and prepared in house and previously used in 2) and embryos returned to the incubator. Day 6 embryos produced at UF were shipped overnight at 38.5°C in serum supplemented culture medium to MU. On day seven, blastocyst-stage IVP embryos were selected, washed in BioLife Holding & Transfer Medium (AgTech; Manhattan, KS), and loaded in groups of two into 0.25 cc yellow, direct transfer and irradiated straws (AgTech) and kept in a Styrofoam box until ET. Blastocysts were transferred to synchronized recipient females on day seven after GnRH injection Maternal blood was collected on D55 and D105 of gestation. CIDR: controlled internal drug release, an intravaginal progesterone releasing device. PG: prostaglandin. GnRH: gonadotropin releasing hormone. AI: artificial insemination. IVF: *in vitro* fertilization. D0: day of AI or IVF.

**Supplemental Figure 2.**
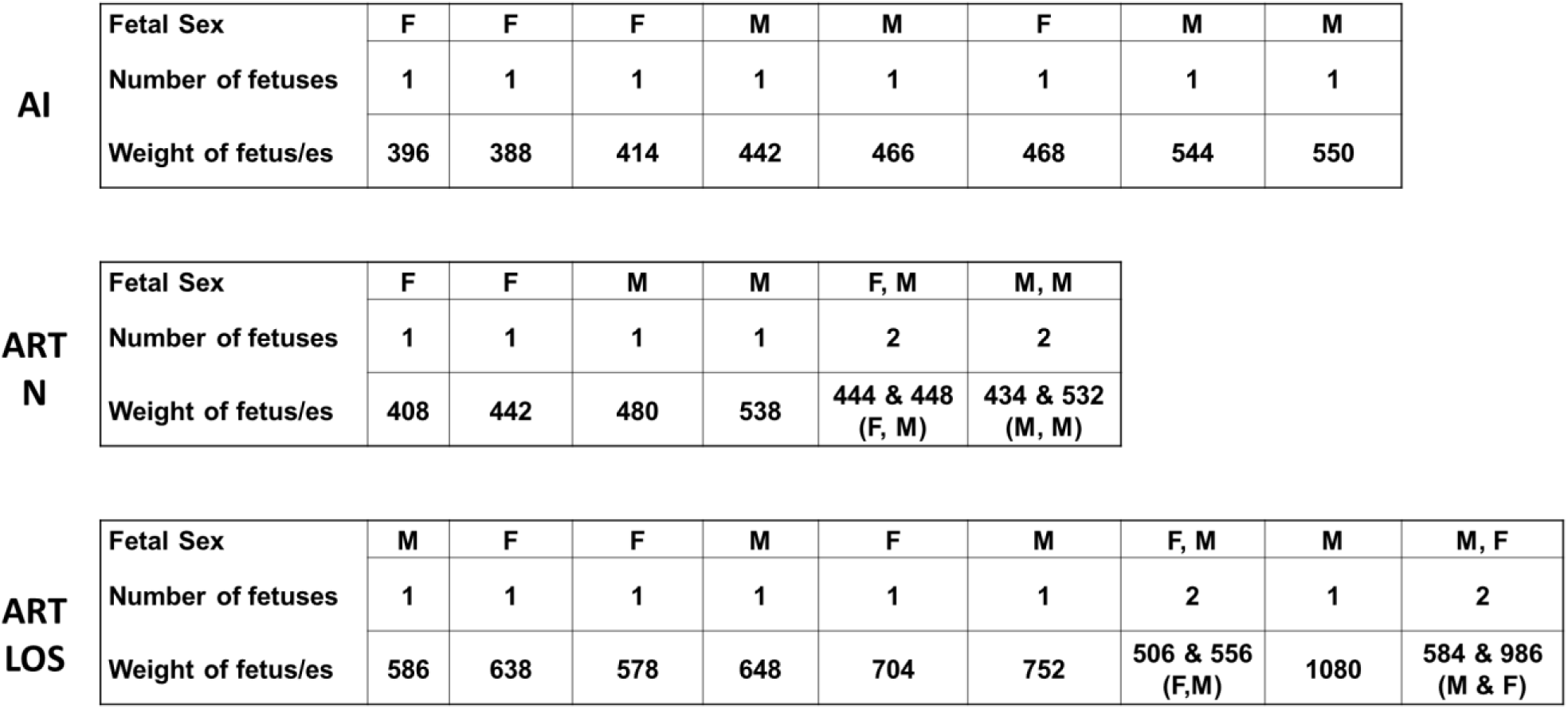
Information on the dams used for WBC transcriptome analyses. AI = artificial insemination (i.e. control). ART-N = embryos were produced by *in vitro* procedures and were <97% of the control’s weight at D105. ART-LOS embryos were produced by in vitro procedures and were ≥97% of the control’s weight at D105. F = female. M = male.

**Supplemental Figure 3.**
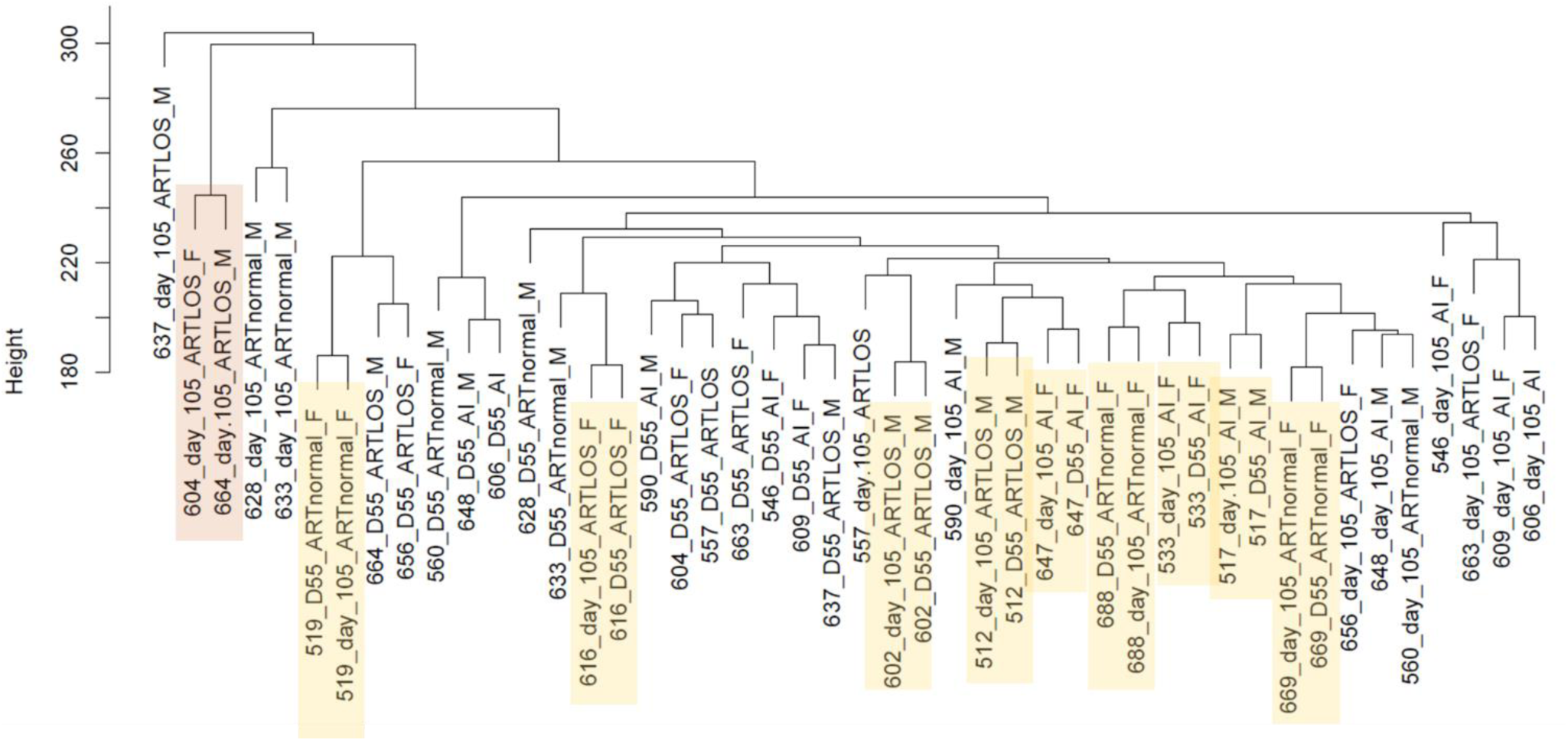
Unsupervised Hierarchical Clustering. Samples highlighted in orange are the WBC transcriptomes of the females carrying the two largest LOS (604 and 664). Samples highlighted in yellow clustered by female irrespective of the blood having been collected on D55 and D105 of pregnancy and during winter and summer respectively. AI = artificial insemination (i.e. control). ARTnormal = embryos were produced by *in vitro* procedures and were <97% of the control’s weight at D105. ART-LOS embryos were produced by in vitro procedures and were ≥97% of the control’s weight at D105. F = female. M = male.

**Supplemental Figure 4.**
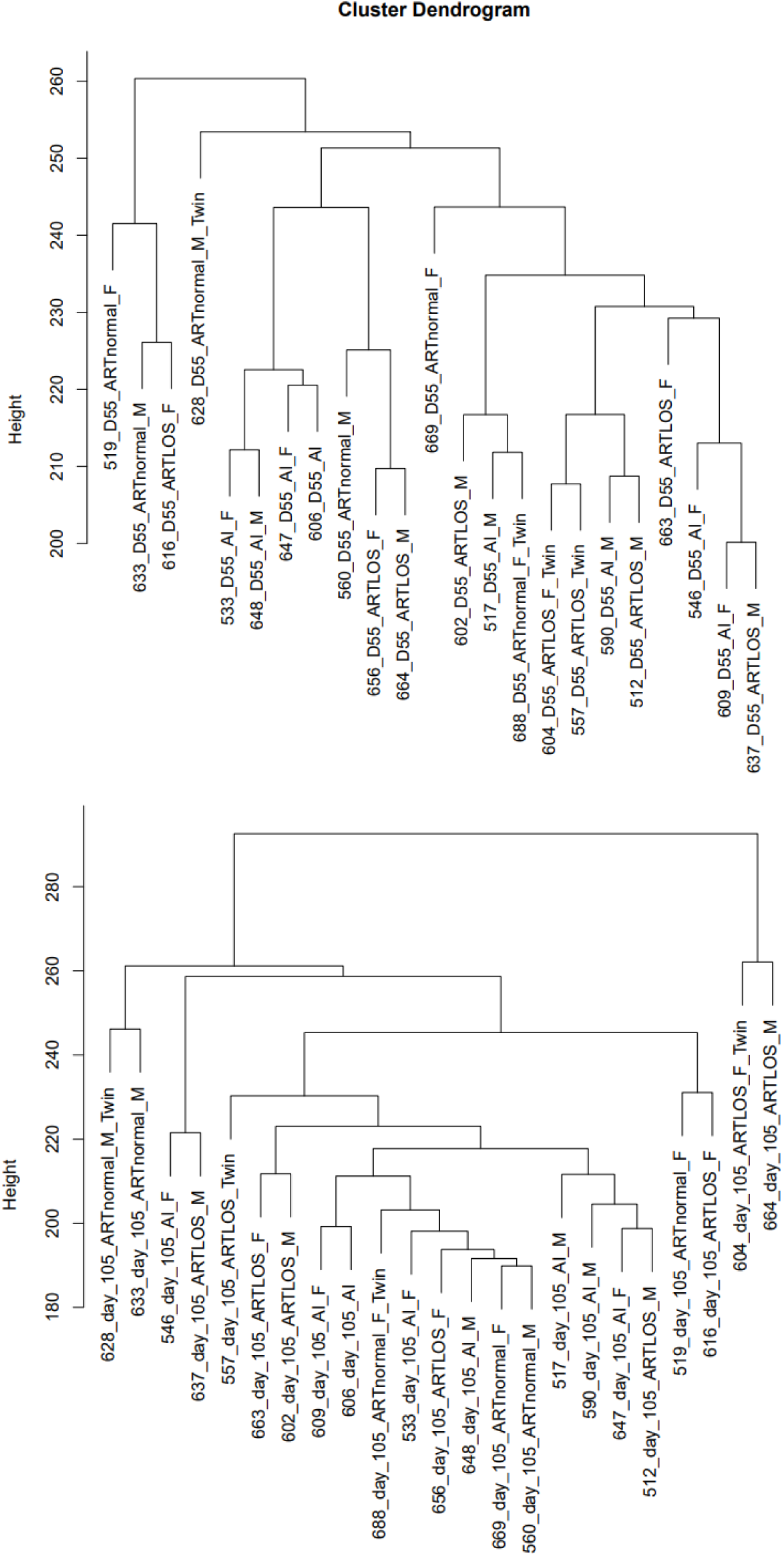
Unsupervised Hierarchical Clustering for D55 and D105 samples. Labels as above.

